# Electrophoresis-Correlative Ion Mobility Deepens Single-cell Proteomics in Capillary Electrophoresis Mass Spectrometry

**DOI:** 10.1101/2024.09.11.612533

**Authors:** Bowen Shen, Fei Zhou, Peter Nemes

**Affiliations:** Department of Chemistry & Biochemistry, University of Maryland, College Park, MD 20742

**Keywords:** Single cell, capillary electrophoresis, mass spectrometry, proteomics, *Xenopus laevis*

## Abstract

Detection of trace-sensitive signals is a current challenge is single-cell mass spectrometry (MS) proteomics. Separation prior to detection improves the fidelity and depth of proteome identification and quantification. We recently recognized capillary electrophoresis (CE) electrospray ionization (ESI) for ordering peptides into mass-to-charge (m/z)-dependent series, introducing electrophoresis-correlative (Eco) data-independent acquisition. Here, we demonstrate that these correlations based on electrophoretic mobility (µ_ef_) in the liquid phase are transferred into the gas phase, essentially temporally ordering the peptide ions into charge-dependent ion mobility (IM, 1/K_0_) trends (ρ > 0.97). Rather than sampling the entire IM region broadly, we pursued these predictable correlations to schedule narrower frames. Compared to classical ddaPASEF, Eco-framing significantly enhanced the resolution of IM on a trapped ion mobility mass spectrometer (timsTOF PRO). This approach returned ∼50% more proteins from HeLa proteome digests approximating to one-to-two cells, identifying ∼962 proteins from ∼200 pg in <20 min of effective electrophoresis, without match-between-runs. As a proof of principle, we deployed Eco-ddaPASEF on 1,157 proteins by analyzing <4% of the total proteome in single, yolk-laden embryonic stem cells (∼80-µm) that were isolated from the animal cap of the South African clawed frog (*Xenopus laevis*). Quantitative profiling of 9 different blastomeres revealed detectable differences among these cells, which are normally fated to form the ectoderm but retain pluripotentiality. Eco-framing effectively deepens the proteome sensitivity in IMS using ddaPASEF, raising the possibility of a proteome-driven classification of embryonic cell differentiation.

## INTRODUCTION

During healthy development, the fertilized egg (zygote) must divide to establish complex cell heterogeneity to pattern the vertebrate body plan in space and time. Knowledge of the proteome, the full set of proteins within a cell, is critical to understanding this process and controlling it on demand. Since the inception of discovery single-cell MS proteomics using the South African clawed frog (*Xenopus laevis*),^1–3^ incredible research progress has been made to uncover proteomic differences among various cell types, focusing recently also on stem cells.

Blastomeres were measured in embryos of the *X. laevis*^1–7^, zebrafish^6^, and mouse^8–9^ as well as oocytes of the mouse^9^ and recently also human volunteers^10^ (partially reviewed in ^11–15^). These studies uncovered many challenges relating to MS proteomics on single stem cells. Blastomeres differentiating in live embryos are “notorious” for shrinking rapidly in size and proteome content upon each cell division and move to complex spatiotemporal locations as tissues are formed. In *X. laevis*, a popular model in health and disease research, the presence of abundant yolk proteome (e.g., ∼90% vitellogenin) complicates matters.^16–17^ There is a high and still unmet need to develop validated next-generation technologies to advance MS proteomics on embryonic stem cells at all stages of vertebrate development, including gastrulation when morphogenic movements begin to set up the embryonic body plan.

At present, smart data acquisition is the focus of active research to deepen the detectable portion of the single-cell proteome. Early on, capillary electrophoresis (CE) ESI was tailored to data-dependent acquisition, identifying up to ∼400 proteins from ∼10 ng of proteome from giant *X. laevis* blastomeres on a time-of-flight (TOF)^3^ and ∼800 on an orbitrap mass spectrometer (Q-Exactive Plus)^2, 6^. Meanwhile the frog also helped adopt isobaric tags to take a leap in the thoughput and accuracy of quantification in single-cell proteomics^3^. Liquid chromatography (LC) reported ∼650^4^ to ∼3,500^7^ proteins from a cell from the 50-cell *X. leavis* embryo, analyzing ∼200 ng proteome. Data-independent acquisition (DIA) expanded LC-MS to ∼1,600 proteins from 40 ng of proteome amount.^5^ SCoPE2 leveraged isobaric quantification to enhance detection sensitivity by introducing a carrier channel.^18^ Recently, DIA and wide-window acquisition (WWA) expanded the sensitivity of data-dependent acquisition DDA to single-cell studies.^19–21^ Real-time search-assisted acquisition, such as RETICLE^22^ and pre-defined inclusion list of peptides of interest (pSCoPE^23^), doubled the coverage to ∼1,400 proteins from single mammalian cells. High-field asymmetric ion mobility spectrometry (FAIMS) reduced isobaric interferences, identifying ∼1,000 proteins in mammalian cell.^24^ Trapped ion mobility time-of-flight (timsTOF) MS with parallel accumulation-serial fragmentation^25^ measured 2,000 proteins in sorted cells.^26^ Most recently, a yolk-depleted carrier was developed to improve MS signal selection despite high proteome interference, returning 15-times more proteins from single blastomeres in *X. laevis.*^17^

Capillary electrophoresis (CE) has been gaining momentum as an alternative to chromatography for trace-sensitive MS, particularly single-cell proteomics. Separation based on electrophoresis is highly efficient and scalable to single cells.^15, 27^ We^2, 6, 28^ and others^29–30^ used CE-MS to analyze picograms^31^ to tens of nanograms^27^ of proteomes from single cells. Using classical DDA, up to ∼1,200 proteins were measured among dissected blastomeres^2–4, 7^ as well as *in situ*^6^ or *in vivo*^28^ in the live *X. laevis* embryo.^6^ DIA expanded CE-MS sensitivity to 1,387 protein during a 15-min separation window from these embryonic cells.^32^ Ion mobility separation measured 1,894 proteins from 10 ng of neuronal proteome digest on a timsTOF PRO.^33^ CE-ESI was applied to 1,314 proteins from 1 ng of the HeLa proteome digest using high-field asymmetric ion mobility spectrometry (FAIMS).^34^ On-capillary cell lysis found 17-to-40 proteins and 23-to-50 proteoforms in single HeLa cells,^29^ while the spray-capillary CE-MS platform reported ∼170– 260 proteoforms from ∼50 cells.^35^ A DDA ladder improved quantification to lower-abundance proteins, reporting ∼165 proteins from ∼150 pg of somal proteome.^36^ Recently, we introduced the concept of electrophoresis-correlative (Eco) ion sorting, where charge-dependent m/z ordering of peptides during electrophoresis deepened the proteome coverage by ∼48% in DIA on an orbitrap mass spectrometer (Q-Exactive Plus, Thermo Fisher Scientific).^37–38^

Here, we introduce the concept of Eco-driven IMS (Eco-IMS) to profile the proteomic state of single blastomeres in the animal cap, a tissue of pluripotent cells, in the vertebrate (*X. laevis*) embryo. This report follows the original disclosure of the technology at the 2023 annual meeting of the American Society for Mass Spectrometry.^37^ With an ability to impart little internal energy to molecules during ionization^39–40^, ESI is broadly used for analyzing the liquid phase via detecting the chemistry of the ions that are generated in the gas phase. We reasoned that CE and IMS perform separation using similar principles: charged species are moved in an electric field. Therefore, we proposed that Eco-sorting based on charge (z), mass (m), and mass-to-charge ratio (m/z) during CE-ESI^2, 37^ would be preserved into the gas-phase, essentially sorting peptide ions based on similar metrics and their resulting ion mobility. To thest this hypothesis, we set out to correlate the detected migration time (MT), m/z, µ_ef_, and K_0_. Next, we benchmark the sensitivity and reproducibility of Eco-IMS for proteome identification and quantification on single-cell equivalent (∼200 pg) amounts of HeLa proteome digests. As a proof of principle, we deploy Eco-IMS to profile blastomeres that form the animal cap in the *Xenopus laevis* blastula. Using unsupervised multivariate and statistical data analysis tools, we systematically quantify the proteomic state of 9 cells in pursuit of cell heterogeneity in this pluripotent cell population critical to vertebrate development.

## MATERIALS AND METHODS

### Materials

The following solvents and chemicals were HPLC-grade and from Thermo Fisher Scientific: acetic acid (AcOH), acetonitrile (ACN), formic acid (FA), and methanol (MeOH). Ammonium bicarbonate (AmBic) was obtained from Avantor (Center Valley, PA). Thermo Fisher Scientific supplied the HeLa protein digest standard (part no. 88329, Pierce, Rockford, IL) and N-dodecyl-β-D-maltoside (DDM, part no. 89903, Rockford, IL). The ESI emitter in the CE-ESI setup was fabricated from a pulled borosilicate glass capillary (0.75/1.00 mm inner/outer diameter, part no. B100-75-10, Sutter Instrument, Novato, CA). The fused silica capillaries for CE were obtained from Polymicro Technologies (40/105 µm inner/outer diameter, part no. 1068150596, Phoenix, AZ) and used without further modification. MS-grade Trypsin Platinum (part no. VA900A, Promega, Madison, WI) was used for proteome digestion. All protein and peptide samples were processed as a microarray of droplets on a fluorosilane-coated plate (termed as “microplate” hereon, Cytonix LLC, Silver Spring, MD).

### Solutions

The *background electrolyte* (BGE) for CE was composed of 1 M FA and 25% (v/v) ACN (resulting pH 2.3). The *CE-ESI sheath solution* contained 10% (v/v) MeOH and 0.5% (v/v) AcOH. The *sample solvent* was prepared of 50% (v/v) ACN and 0.05% (v/v) FA. For dissociating the embryo to a cell suspension, the Newport buffer was prepared as instructed elsewhere.^41^ The cell *lysis solution* contained 0.2% DDM in 50 mM AmBic.

### Animal Care and Embryology

Sexually mature *Xenopus laevis* frogs were sourced from Nasco (Fort Atkinson, WI) or Xenopus1 (Dexter, MI). All procedures related to the humane care and use of *X. laevis* were authorized by the Institutional Animal Care and Use Committee at the University of Maryland, College Park (Approval nos. R-FEB-21-07 and R-FEB-24-05). To minimize biological variablity, all the embryos in this study were obtained through the natural mating of a single pair of parents. Embryos with stereotypical cleavage were chosen at the two-cell stage and cultured to stage 8 following our earlier studies. Each embryo and cell was labeled with a unique identifier, although this intormation was intentially hidden during data analysis and only revealed to faciliate the interpretation of results.

### Single-Cell Collection and Proteome Processing

Using a micro knife (No. 10318-14, Fine Science Tools, Foster City, CA), the vitelline membrane surrounding the stage-8 embryos was carefully peeled away. As illustrated in **Figure 1**, the tissue covering the animal cap was identified based on pigmentation and location, dissected, and transferred into a Petri-dish filled with the Newport buffer (**Fig. 1**). The tissue was dissociated to a suspension of cells under gentle agitation with a hair loop. The resulting cells were rinsed of salts via quick transfer into HPLC water using a pipette tip, without visual lysis or damage to the cells confirmed under a stereomicroscope. Finally, these blastomeres were individually pipetted into an array on a fluorosilane-coated slide (microplate). Each cell was lysed by manual addition of 500 nL of the *lysis buffer*, with incubation at room temperature for 10 min. The extracted single-cell proteomes were denatured by rapid heating to 60 °C for 15 min, then digested with 500 nL of 0.1 µg/µL trypsin platinum (in 50 mM AmBic buffer) at 40 °C over 1 h. Liquid evaporation was compensated for by addition of 400 nL of HPLC water to each sample every ∼5 min. The resulting single-cell proteome digests were allowed to dry at room temperature, before storage at –80 °C (the “samples”) until analysis.

**Figure 1.**
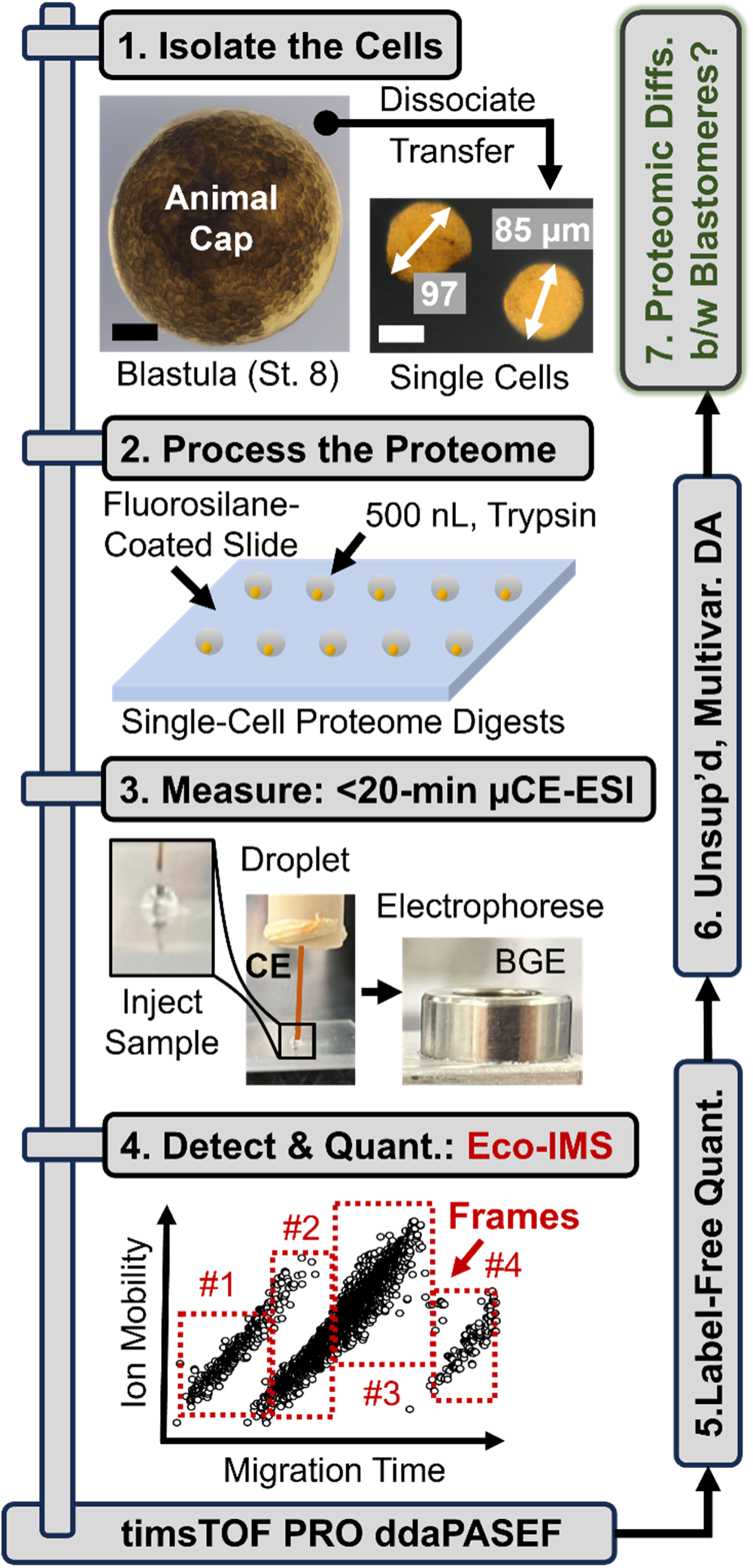
Our experimental approach to develop, test, and validate Eco-IMS for single-cell proteomics. As a proof-of-principle, blastomeres were dissociated from the animal cap of the *X. laevis* gastrula (stage 8, animal cap view shown) and transferred individually onto a fluorosilane-coated plate, where each cell was trypsin-digested in ∼500 nL of droplets (see details in the **Methods**). Ca 4% of the cell proteome was electrophoresed in <25 min on a previously validated µCE-ESI platform.^27, 32^ The generated peptide ions were separated based on ion mobility (IM), viz. collision cross section, before fragmentation via CID under control by ddaPASEF. The abundance of proteins was estimated via label-free quantification (LFQ), feeding quantitative information into an unsupervised data analysis model to seek differences between the blastomeres. Scale bars, 250 µm (black) and 50 µm (white).

### CE-nanoESI-MS

The samples were reconstituted in the sample solvent for analysis using CE-MS. The CE-nanoESI platform was constructed and operated as detailed earlier.^6^ In this study, the proteome digest was electrophoresed in a 100-cm capillary by applying +23 kV (vs. Earth ground) to the BGE filling the inlet end of the capillary. The outlet end of the CE capillary was connected to a CE-nanoflow ESI interface that was electrokinetically pumped at 700–1,100 V potential. This interface was constructed following earlier designs^42^ and operated in the cone-jet regime^6^ for maximal ionization sensitivity^39^. The emitter tip of the CE-ESI source was positioned ∼1 mm from the orifice of the mass spectrometer. A high-resolution camera, equipped with a long-working distance objective, was used to confirm the spraying regime as per our pevious studies.^2, 39^ The generated peptide ions were detected on a timsTOF PRO mass spectrometer (Bruker Daltonics, Billerica, MA) executing the ddaPASEF under the classical or Eco control (details below). The conditions of MS detection were optimized to the type of proteome that was measured, as follows.

#### MS Settings for the HeLa Proteome Digest

Eco-IMS was developed on 500 pg portions of the HeLa proteome digest (approximating to ∼2 cells), before testing performance on ∼200 pg of proteome amounts, or ca. a single cell. In the classical IMS, ions were analyzed between 1/K_0_ = 0.60–1.60 over the entire duration of separation. In Eco-IMS, this range of IM was sliced into 4 different frames following optimization (Frame #, separation time [min], 1/K_0_ [V s cm^-2^]): Frame #1, 20–27, 0.65–1.00; #2, 27–32; #3, 32–40, 0.80–1.40; and #4, 40–45, 0.70–1.05. The following parameters were set global across this study, irrespective of the method of data-acquisition method used: MS^1^ scan range: *m/z* 300–1,500; ramp times, 100 ms for 500 pg and 200 ms for 200 pg of the HeLa proteome digest; number of PASEF scans, 12 for 500 pg and 6 for 200 pg of the HeLa proteome digest; target intensity, 10,000 counts; and threshold, 1,500 counts; high-sensitivity detection mode, enabled; dynamic exclusion, 0.2 min. Only ions with +2 and +3 charges were selected for detection in this study. All other parameters were set to the default.

#### MS Settings for Single Embryonic Cells

Each *X. laevis* single-cell proteome digest was reconstituted in 400 nL of the *sample solvent*. Ca. 15 nL (∼2 ng) of each single-cell proteome were loaded from the fluorosilane-coated microplate into the CE separation capillary. In the classical IMS approach, 1/K_0_ was scanned between 0.6–1.6 for the entire duration of separation. In Eco-IMS, the following 4 frames were implemented following optimization (Frame #, times of duration [min], 1/K_0_ [V s cm^-2^]): #1, 0–30, 0.6–1.1; #2, 30–38, 0.6–1.4; #3, 38–44, 0.75–1.3; and #4, 44–50, 1.0–1.4. All other parameters were set the same as for the HeLa standards.

### Data Analysis

The primary MS–MS/MS files were processed with MSFragger 20.0^43^. Proteins were identified against the Human proteome (UP000005640, downloaded from UniProt on July 9^th^, 2023, containing 20,523 entries) or *Xenopus* proteome (UP000694892, downloaded from UniProt on Jan 10^th^, 2022, containing 43,236 entries). The search parameters were the following: static modification, cysteine carbamidomethylation; dynamic modifications, methionine oxidation; minimum peptide length, 5 amino acids; maximum peptide length, 50 amino acids; and maximal missed cleavage sites, 2. Technical replicates of each sample were analyzed together with the match-between-runs (MBR) function enabled. All other parameters were set to the default.

### Scientific Rigor

The HeLa proteome digest was measured in 3–5 technical replicates (same sample analyzed repeatedly). In this study, n = 9 different single blastomeres were isolated from embryos that were obtained from the natural mating of the same pair of *X. laevis.* The cells were isolated and measured in a randomized order. The custom-built CE-ESI platform was validated on the timsTOF PRO mass spectrometer daily using the HeLa reference proteome digest.

### Safety

General safety measures were followed when working with chemicals and biological specimens. Capillaries and ESI emitters were carefully handled to minimize puncture hazard. All electrically conductive elements of the CE-ESI platform were shielded or grounded (to true Earth). Its protective enclosure with a safety interlock provided additional safety mesure from an accidental electroshock.

## RESULTS AND DISCUSSION

### Wet Ion Sorting Goes Dry

Our experimental design to developing Eco-IMS is presented in **Figure 1**. The approach was refined, then validated on single-cell-equivalent amounts of HeLa proteome digests (∼200 pg). We demonstrated Eco-IMS for the protomic profiling of single blastomeres in the mid-stage *X. laevis* embryo (Nieuwkoop-Faber stage 8). The animal cap was selected as model tissue in this study, as this layer of stem-like cells are fated to give rise to the embryonic ectoderm, yet remain susceptible to induction to different fates (reviewed in ^44^). The proteome of each of n = 9 different blastomeres were extracted and digested in 500 nL droplets of trypsin in 50 mM AmBic, directly on a fluorosilane-coated glass slide. These resulting digests were analyzed on a custom-built µCE-ESI platform as per our protocols.^2, 27–28^ The generated ions were measured on a timsTOF PRO mass spectrometer. Detection was executed under ddaPASEF control, selected to obtain 100% duty cycle during tandem MS.^25^ The proteome profile was approximated by calculating the label-free quantification (LFQ) index of each protein. The resulting cell profiles were systematically compared through an unsupervised multivariate–statistical analysis. Discovery single-cell proteomics raised a potential to appreciate molecular differences between these pluripotent stem-like cell population.

Our analytical hypothesis was empirically testable. We reasoned that separation based on electrophoretic mobility (µ_ef_) differences in the liquid phase during CE share mechanistic similarities to IMS leveraging ion mobility differences in the gas phase—that is the movement of a charged species (in a media) under an external electric field. **Figure 2** tests the theory through interconnected data analyses of 500 pg of the HeLa proteome digest standard, approximating to ca. 2 cells. **Figure 2A** exemplifies electrophoretic sorting of the peptides into charge- and m/z-dependent trends, the basis of Eco-MS^37^. The recorded MTs were in exquisite correlation with the theoretical mobilities that we calculated based on the experimental peptide sequences (**Fig. 2B**) following an earlier model^45^. This allowed us to ascribe the principal mechanism of separation to be electrophoretic in nature in our CE system (**Fig. 2B**). This knowledge was logically necessary to explore a relationship between the liquid-phase µ_ef_ and gas-phase ion mobilities (1/K_0_). High Pearson correlation moments (ρ = 0.81–0.97) across the charge states validated a general correlation between the separation mechanisms (**Fig. 2C**). This fundamental relationship in turn raised the potential to transfer CE-sorting to the domain of IM. Indeed, the measured 1/K_0_ values exhibited high correlation with MTs (ρ > 0.91, Pearson correlation), as is shown in **Figure 2D**. This result validated the working hypothesis of this study.

**Figure 2.**
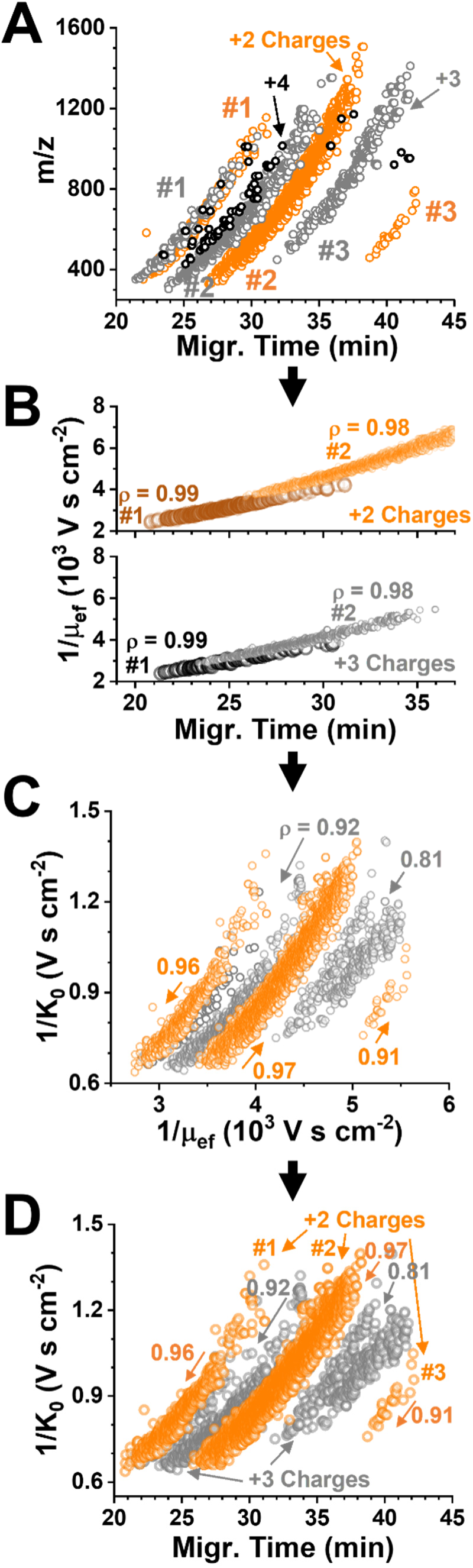
Experimental validation of electrophoretic ion sorting for ion mobility mass spectrometry (IMS). **(A)** Representative relationship between the detected peptide m/z and CE migration time. **(B)** Exquisite correlation between the electrophoretic mobility (µ_ef_) and migration time regardless of charge state, revealing ion sorting based on mobility in the liquid phase. **(C)** Correlations were strong-to-excellent between the measured gas-phase ion mobility (1/K_0_) and liquid-phase µ_ef_, suggesting Eco-sorting also along the IM domain. **(D)** Indeed, CE was found to essentially order the peptides into charge-dependent gas-phase IM trends. We leveraged these correlations to improve the utilization of IM bandwidth via framing along the separation time. Key to all panels: Pearson correlation moment, ρ.

This correlation between ion sorting practically “wet” and “dry” served the basis for developing Eco-IMS in this work. We recently found MT-dependent scheduling of the m/z domain to beenfit sensitivity and quantification.^38^ This led to our reasoning that MT-dependent framing would also advance detection along the domain of IM separation. Compared to the classical approach to scan the entire range of IM, we leveraged Eco-sorting to analyze only the ranges of IM that peptides actually occupied with separation. This in turn promised to improve ion accumulation for higher sensitivity while spare detection bandwidth for MS/MS. **Figure 3A** presents the approach, showcasing only the +2 charges for simplicity. In agreement with our previous findings,^37–38^ this charge state accounted for ∼86% of all identifications and were sufficient for this study; the other charge states can, of course, (also) be included in future embbodiments to deepen sensitivity. We scheduled Eco-IMS to temporally schedule ddaPASEF along the following 4 frames tailoring to these +2 states (Frame no., ranges of separation time [min], min/ion mobility [V s cm^-2^]): #1, 20–27, 0.65–1.00; #2, 27–32, 0.60–1.20; #3, 32–40, 0.80–1.40; and #4, 40–45, 0.70–1.05. To balance technical complexity, Frame #2 captured two apparent subframes with an intentionally broader range of IM. As the benchmark of performance, the classical ddaPASEF was performed with the default parameters: the IM was sampled between the entire 0.6–1.6 V s cm^-2^ range with separation.

**Figure 3.**
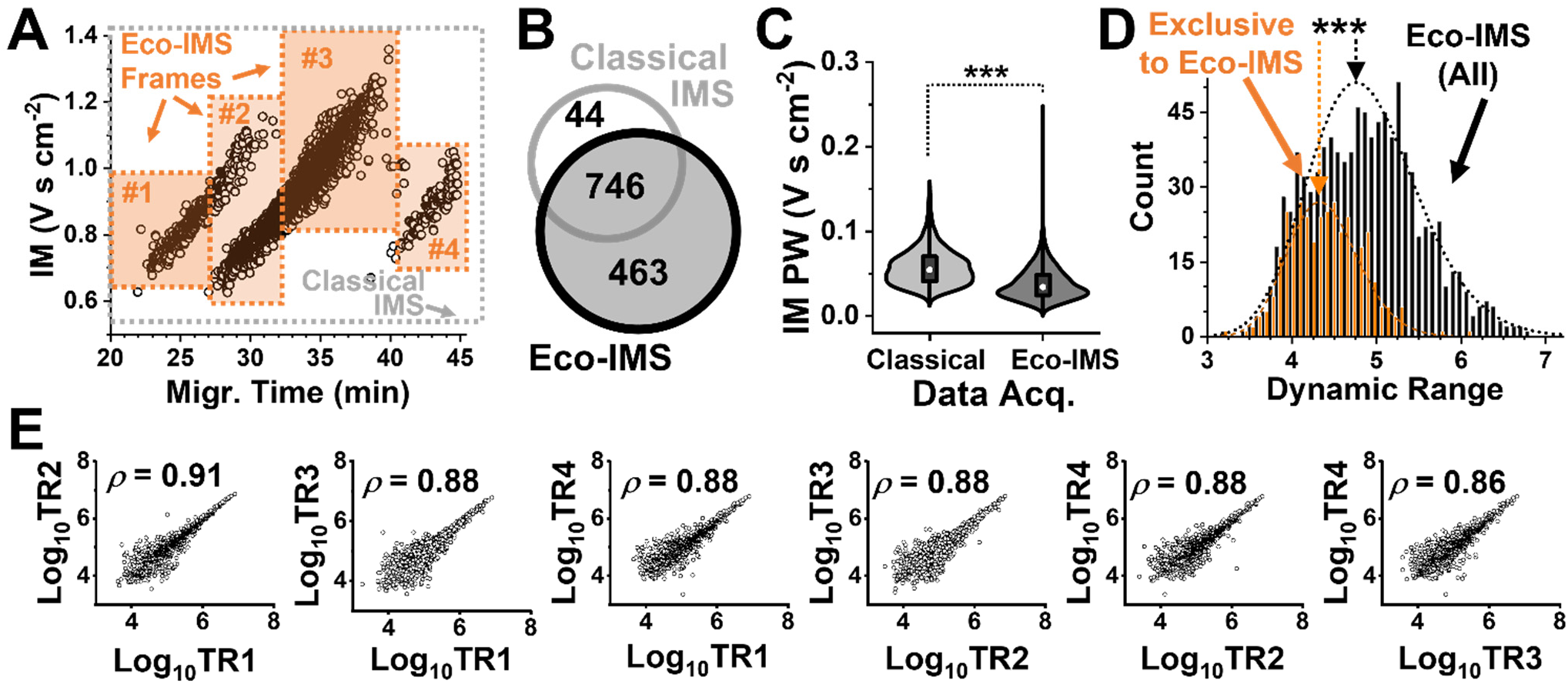
Development of Eco-IMS and performance characterization. Ca. 500 pg of the HeLa proteome digest, approximating to 2 cells, were measured on a timsTOF PRO executing ddaPASEF. **(A)** We developed Eco-IMS by scheduling IM scans according to the ordered separation of the peptides. The example shows configuration of the method based on 4 frames (labels #). For simplicity, Frame #2 was designed to capture the overlapping tailes of #1 and #3. Benchmarking of **(B)** proteome identification and **(C)** peak widths (PW) during ion mobility (IM) as well as **(D)** the dynamic range and **(E)** reproducibility of quantification among 4 technical replicates (TR1–4). Protein concentrations were estimated based on label-free quantification (LFQ) without match-between runs. Key: ***, *p* < 0.001 (Mann-Whitney U test).

We benchmarked the performance of proteome identification. Using identical cycle time, we projected narrower spans of IM to allow for slowing down the rate of gas-phase separation in Eco-IMS, therefore improving IM resolution. We anticipated other metrics building on resolution to also improve, including detection sensitivity. Indeed, Eco-IMS identified 1,209 HeLa proteins, marking a 53% improvement from the 790 proteins detected under the classical approach (**Fig. 3B**). Ca. 94% of the proteins that were identified by the classical ddaPASEF were also measured by the Eco-driven. The electronic **Supplementary Information** lists the proteins that were detected using classical (**Table S1**) and Eco (**Table S2**) IMS. As revealed in **Figure 3C**, the meadian width of the recorded IM peaks was significantly lower in the Eco-IMS dataset than the classical (*p* < 0.001, Mann-Whitney U test). Therefore, narrower IM frames via Eco-MS improveded the resolution of IM separation and sensitivity of proteome detection.

We then investigated the impact Eco-IMS had on quantification. We inquired about the respective concentration of the detected proteins using LFQ. An ∼500 pg portion of the HeLa proteome digest was masured with both methods, each in 4 TRs. The calculated LFQ values were log10-transformed and median normalized (**Methods**). The protein abundances were found to correlate among the TRs and technologies with a Pearson product moment of >0.8 (**Fig. S1**). As shown in **Figure 3C**, proteins that were exclusively quantified by Eco-IMS populated the lower concentration range of the measured proteome (*p* = 3.3 × 10^-41^, Mann-Whitney U test). For the TRs using Eco-IMS, Pearson correlations were high, ranging between 0.88 and 0.91 (**Fig. 3E**). Therefore, Eco-IMS improved the sensitivity of proteome quantification.

We tested the scalability of this sensitivity to single-cell-equivalent proteomes. A 200 pg portion of the HeLa proteome digest was analyzed using Eco-IMS. For such limited proteomes, the spectra quality was enhanced by elevating the accumulation/ramp time to 200 ms.^25^ To maintain a similar duty cycle time, the number of PASEF scans was compensated from 12 to 6. After this optimization, the classical ddaPASEF identified 505, 529, 488, and 506 proteins, accumulating to 661 proteins without MBR (**Table S3**). In contrast, Eco-framing improved identifications to 675, 667, 726, and 680 proteins, amounting to 962 proteins without MBR (**Table S4**). Ca. ∼94% of the identifications from the control were also shared by Eco-IMS (**Fig. 4A**). Gene ontology (GO) annotation (PANTHER version 18.0^46^) revealed appreciable improvements in the coverage of molecular function (**Fig. 4B**) and biological process (**Fig. 4C**) using Eco-IMS. Several important proteins were only detectable using Eco-IMS, including the following: Tubb1, Tubb8, Tubal3 have roles in cell division^47^; the proteins Mcm7, Mcm5, Mcm3, Mcm4, Top2a participate in DNA metabolism; the proteins Eif3l, Eif3cl, Eif3d, Eif3e, Eif3f, and Eif3i are release factors during gene translation.

**Figure 4.**
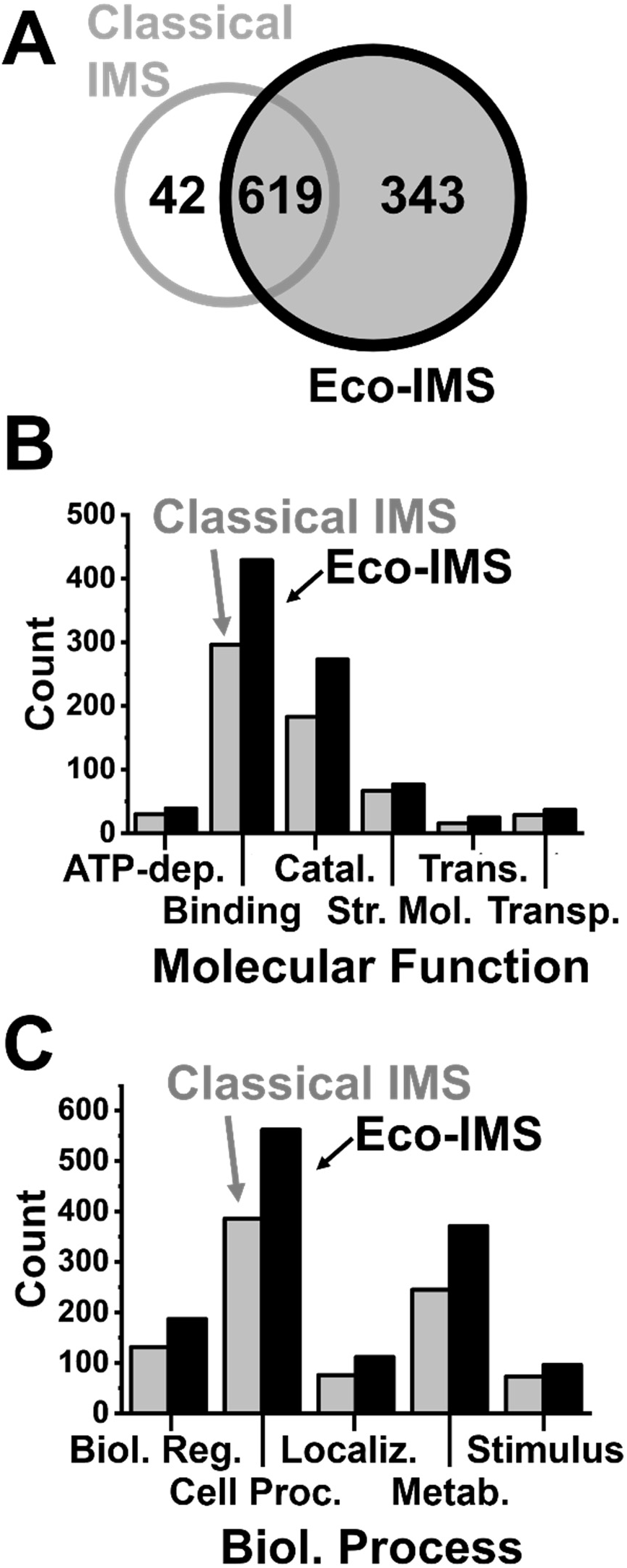
Assessment of scalability to single-HeLa-cell equivalent proteomes. A ca. 200 pg of the standard proteome digest was bechmarked using Eco-IMS against the classical method. **(A)** Comparison of proteome sensitivity, **(B)** molecular function, and **(C)** biological processes based on the annotation of canonical knowledge on the identified proteins. Eco-framing improved the sensitivity of CE-ESI-MS sensitivity to single-cell equivalent proteome amounts.

### Eco-IMS for Profiling a Pluripotent Stem-Cell Population

At this stage, we considered Eco-IMS validated, ready to trace the proteomic state of single cells undergoing differentiation in the *X. laevis* embryo (**Fig. 1**). Earlier studies from our group^2, 6, 27–28^ and others^4–5, 7^ analyzed large cells from this model (100–500 µm in diameter). In this work, we focused on the animal cap, a tissue consisting of pluripotent stem-like cells. As demonsrated in **Figure 5A**, we dissected this tissue from stage 8 embryos and dissociated it to manually isolate n = 9 blastomeres (**Methods**). Each cell was assigned a unique identifier, although this information was only released at the end of the study to aid results interpretation. With an ∼75–100 µm diameter, these cells encompassed only 200–500 pL of volume, containing approximately 35–90 ng of total proteome.^2^ To explore analytical sensitivity toward measuring limited proteomes in future work, we set out to analyze ∼2 ng, or <4% of the whole single-cell proteomes.

**Figure 5.**
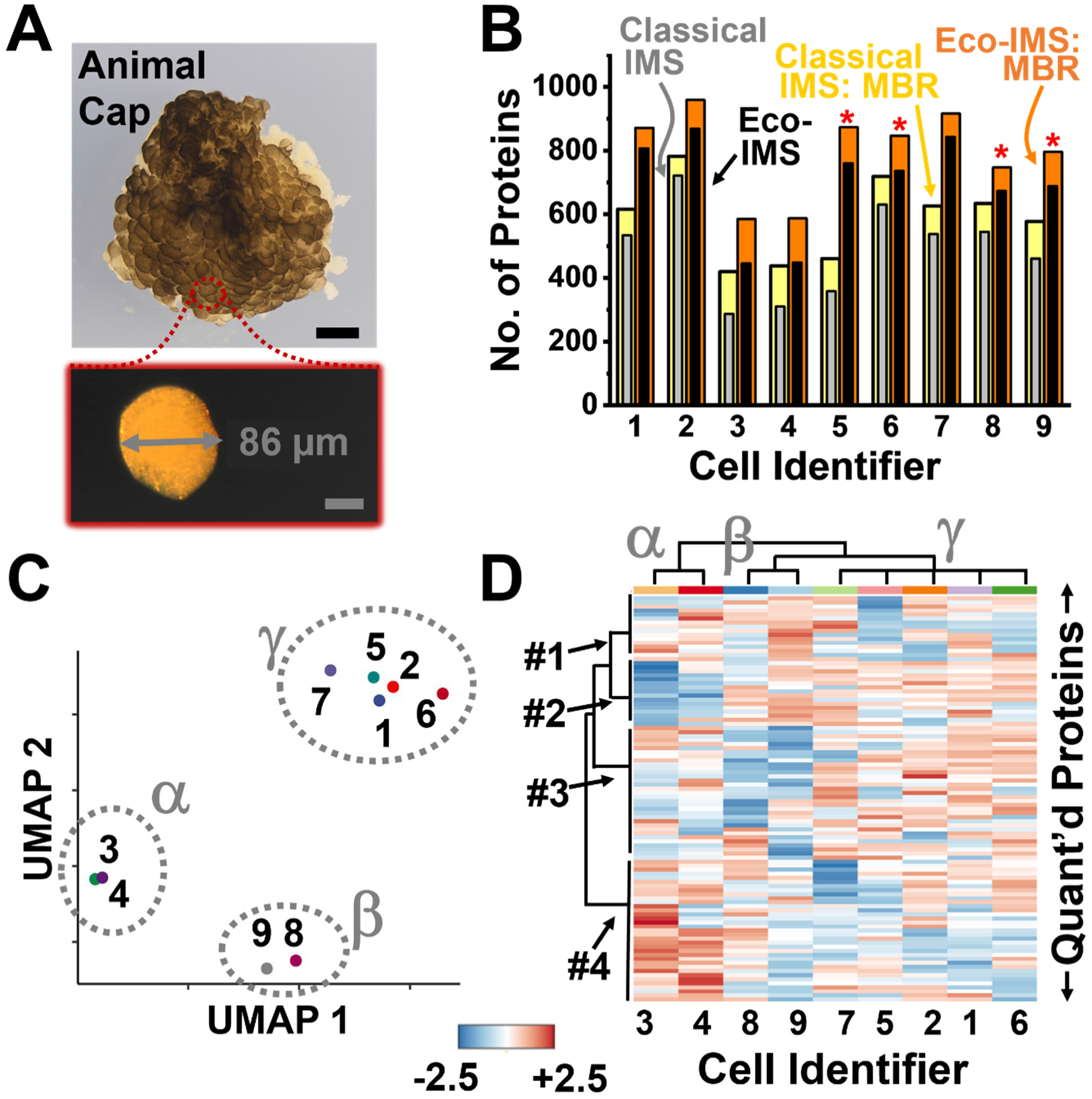
Eco-IMS profiling of stem cell proteomes in the *X. laevis* embryo. **(A)** The animal cap was dissected at mid-blastula stage 8 (top panel) and dissociated to isolate n = 9 different single blastomeres (bottom panel). Scale bars, 125 µm (black) and 50 µm (gray). **(B)** Comparison of proteome coverage from the cells using the classical and Eco-driven IMS on a timsTOF PRO executing ddaPASEF. The results are shown also using match-between-runs (MBR) enabled in MSFragger. **(C)** UMAP profiling of the single cells based on label-free quantification organized the cells into 3 clusters (labeled α, β, and γ). These groups did not correlate with the identity of each embryonic animap cap (labeled 1–9), suggesting that the analysis revealed innate proteomic differences. **(D**) Hierarchical cluster-heatmap analysis on the top 100 most significantly differently quantified (quant’d) proteins revealed systematic differences in protein expression (labels #1–4). The samples were grouped into the same α, β, and γ clusters as on UMAP. Scale bar, z scale. Combined, these results demonstrated the ability of Eco-IMS to survey proteomic differences between embryonic stem cells undergoing differentiation.

Our workflow was modified to adapt flat substrates to process these limited proteomes. Up to this work, our protocols processed giant blastomeres in plastic vials (Eppendorf Lo-Bind) and used manual pipetting to transfer a portion of the sample for CE-MS analysis.^6^ To reduce analyte losses due to nonspecific adsorption on contact sufaces, we processed the single-cell proteomes in droplets. As shown in **Figure 1**, a fluorosilane-coated microplate was employed to reduce contact angles. Replicate analyses on the same sample returned a comparable number of proteins between the classical in-vial and microplate fromats (*p* = 0.70, unpaired student t-test, **Fig. S2A**). The log_10_-transformed and median-normalized LFQ abundances were statistically comparable between the technologies (*p* = 0.85, Kruskal-Wallis test, **Fig. S2B**). These LFQ Pearson correlation moments were >0.9 between randomly selected replicates, demonstrating reproducible and ca. 5-times faster sample processing using the microplate.

Each single blastomere was analyzed using both Eco and classical IMS. As the IM vs. MT profile of the *X. leavis* was slightly different than that of the HeLa reference, we developed Eco-scheduling that specifically tailored to the proteome of the frog (settings in the **Methods**). The quantitative coverage of the proteome was improved by engaging MBR in MSFragger (**Methods**). The results are presented in **Figure 5B**. While classical ddaPASEF measured a total of 909 proteins (**Table S5**), Eco-MS was able to report on 1,157 (**Table S6**). Based on GO analysis (PANTHER version 18.0^46^), 96 of these proteins fulfil roles in biological regulations, 347 proteins partake in metabolic process, and 445 proteins carry out cellular process ranging from cell cycle to division to communication. Among the proteins that were only quantifiable by Eco-MS was also ribosomal protein L4 (Rpl4), which is naturally enriched in the animal cap,^48^ and cell division cycle 42 (Cdc42), a player of cell movement during gastrulation.^49^

The quantitative data also measured gene translation, raising a potential to appreciate differences among the blastomeres. The log10-transformed and median-normalized data were subjected to uniform manifold approximation and projection (UMAP), a popular approach to appreciating cell clusters in multidimensional single-cell data.^50^ As shown in **Figure 5C**, this analysis organized the 9 cell proteomes into 3 clusters. These cells were obtained from the mating of the same one pair of frogs. On revealing the identity of each animal cap, no correlations were found with the groups. Therefore, we ascribe the observed cell heterogeneity to different states of differentiation.

Hierarchical cluster analysis (HCA) was recruited as an orthogonal means to inspect the data. HCA grouped the samples into the same α, β, and γ clusers as UMAP. **Figure 5D** presents the HCA–heatmap based on the 100 most significantly differently quantified proteins (close-up in **Fig. S3**). The dendrograms revealed systematic differences among the relative proteome abundances. Some were strikingly clear. Proteins in Cluster #2 were least abundant in the cells of group α (cells #3 and #4). Multiple proteins in this group were related to the cell cycle and development, including the microtubule plus-end binding protein EB1(participates in mitotic spindle orientation and cell division^51^), adenylyl cyclase-associated protein (partakes in asymmetric cell division and cytokinesis^52^), and miniature chromosome maintenance 7 (regulates DNA replication licensing complex^53^). Therefore, it is possible that these blastomeres may be at different stages of the cell cycle or division. In contrast, proteins in Cluster #4 had the highest levels in cells α and lowest in γ (cells #1, #2, #5, #6, and #7). Many of these proteins were involved in ribosomal synthesis (rplp0, rplp2, translation initiation factor protein central to translation initiation during embryonic development^54^) or mitochondrial function and energy production (e.g., ndufb9 and ndufb10, slc25a3^55^, atp5pb).^56^ It is plausible that these cells were at different protein synthetic stage, perhaps en route to differentiation from the other cell fates.

## CONCLUSIONS

Ths study developed and validated Eco-IMS for single-cell CE-ESI-MS proeomics. Using a timsTOF PRO mass spectrometer executing ddaPASEF, we found CE-driven Eco-framing of IM separation to improve protein identification and quantification. From single-cell equivalent HeLa proteome digests, Eco-IMS employing 4 IM–MT frames on this previous-generation mass spectrometer returned 962 proteins without MBR. This marked a 50% enhancement in proteome coverage compared to the classical ddaPASEF. There are several technological developmental opportunities to further Eco-MS. For example, automation to process cells would allow for exploring larger cell populations, thus beginning to complement already available single-cell transcriptome maps on embryonic cell differentiation. Analogously to our recent findings in Eco-driven DIA,^37–38, 57^ we anticipate Eco-IMS proteome coverage to improve substantially when using a higher number of IM-MT frames; alternatively, the trends may be traced live.

Eco-IMS was able to find previously unknown proteomic differences between blastomeres in the embryonic animal cap tissue. Eco-IMS on our TimsTOF PRO platform identified 1,157 proteins from n = 9 single blastomeres that were manually isolated from the animal cap that we dissected from the developing *X. laevis* embryo. UMAP and HCA, two orthogonal models of multivariate data analysis, independently organized these cells into the same 3 clusters based on the calculated LFQ profiles. HCA suggested these groups to be driven, in part, by systematic differences in protein components related to the control of the cell cycle, development, protein translation, and energy production.

## Acknowledgment

Parts of this research were supported by the National Institute of General Medical Sciences (award nos. R35GM124755 and R35GM5207292 to P.N.), the National Institute of Aging (award no. R01AG088147 to P.N.), the Arnold and Mabel Beckman Foundation (Beckman Young Investigator Award to P.N.), or the University of Maryland, College Park (Start-Up to P.N.).

## Data Repository

All MS–MS/MS primary files and the MS/MS spectral libraries, and the HeLa and Xenopus proteomes were deposited to the ProteomeExchange Consortium via the PRIDE partner repository with the data set identifier PXD05302.

## Authorship Contributions

P.N. and B.S. designed the study. F.Z. dissected the *Xenopus laevis* animal caps and isolated the single cells. B.S. prepared and measured the proteome digests. B.S. and P.N. analyzed the data, interpreted the results, and wrote the manuscript. All the authors commented on the manuscript.

