## Supplementary material for "Electrophoresis-Correlative Ion Mobility Deepens Single-cell Proteomics in Capillary Electrophoresis Mass Spectrometry": SI Document

### FIGURES

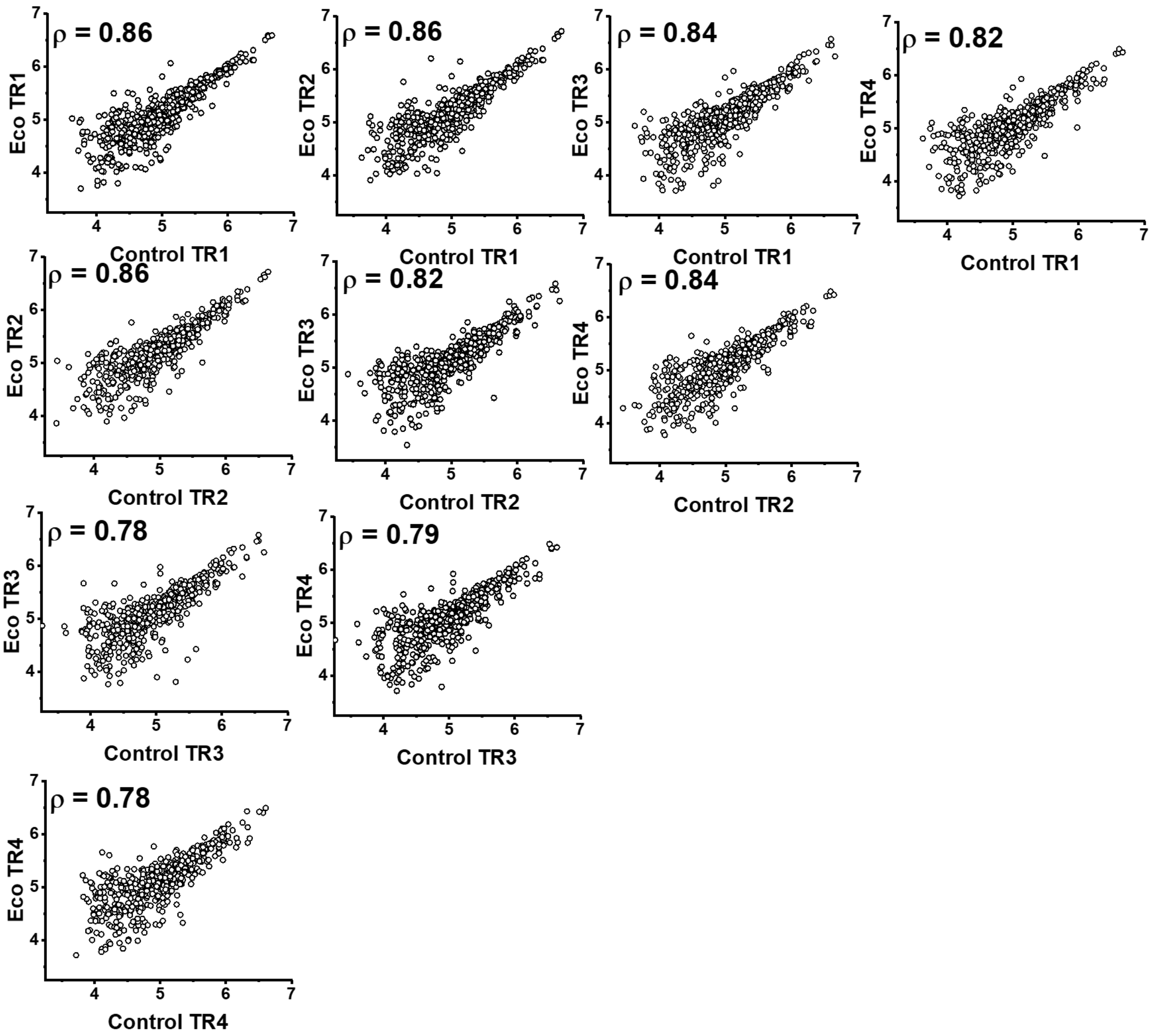

**Figure S1.** Benchmarking of Eco-IMS proteome quantification vs. the classical approach (Control). Ca. 500 pg were measured from the HeLa proteome digest in 4 technical replicates (TR) using each method. Protein concentrations were estimated based on $\log_{10}$-transformed and median-normalized label-free quantification indexes (MSFragger), without using match-between-runs. We considered the resulting Pearson correlation moments of $\rho = 0.78–0.86$ to be sufficient indication of good quantitative fidelity.

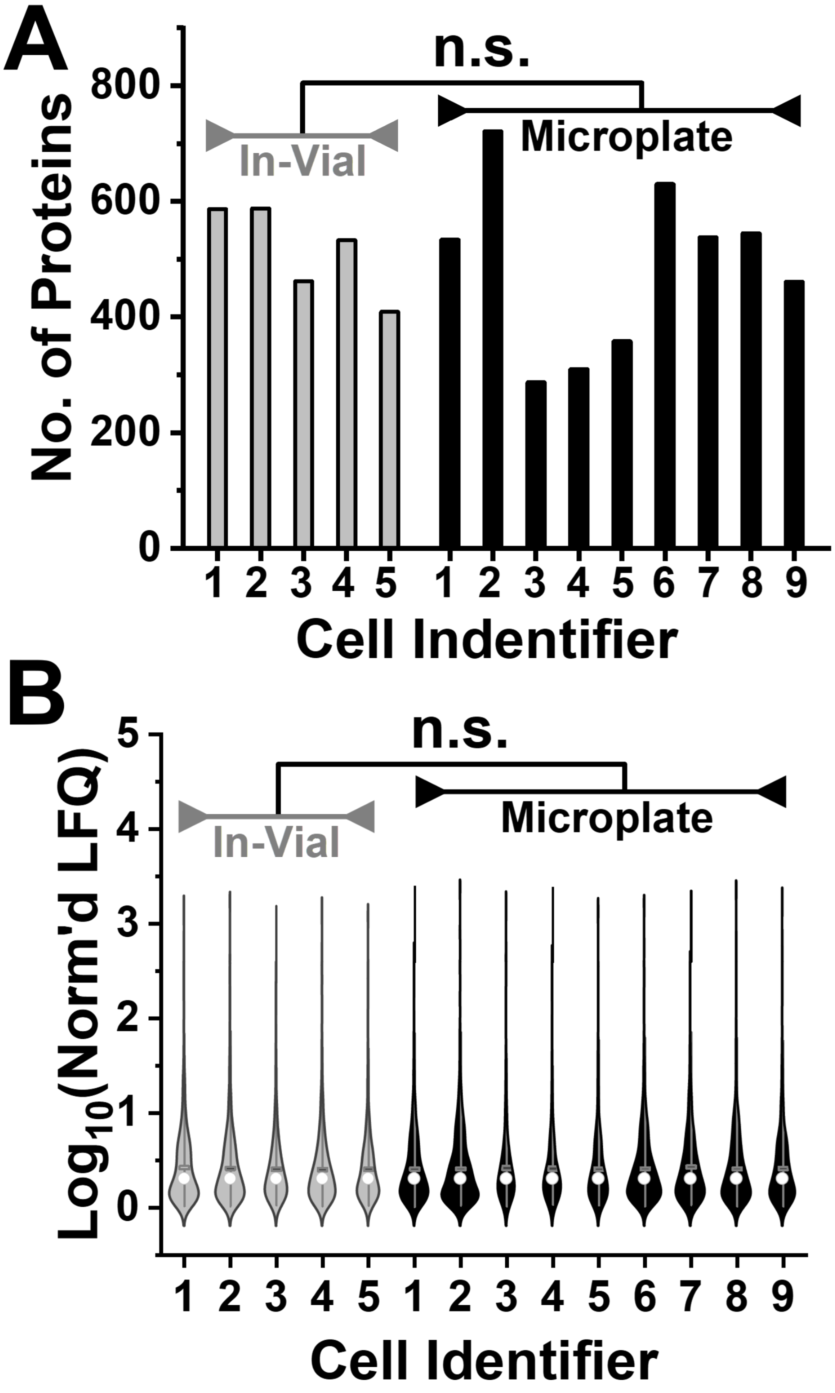

**Figure S2**. Comparison of **(A)** protein identification and **(B)** quantification performance with the sample processing between the classical plastic vials vs. the fluorosilane-coated microplates. The microdroplet format allowed us to digest single-cell proteomes using limited reagents ca. 5-times faster in similar performance between the approaches ($p$ = 0.70). The LFQ data were median normalized and $\log_{10}$ transformed. Key: n.s., not significant (**Fig. S2A**, unpaired student t-test; **Fig. S2B**, Kruskal-Wallis analysis).

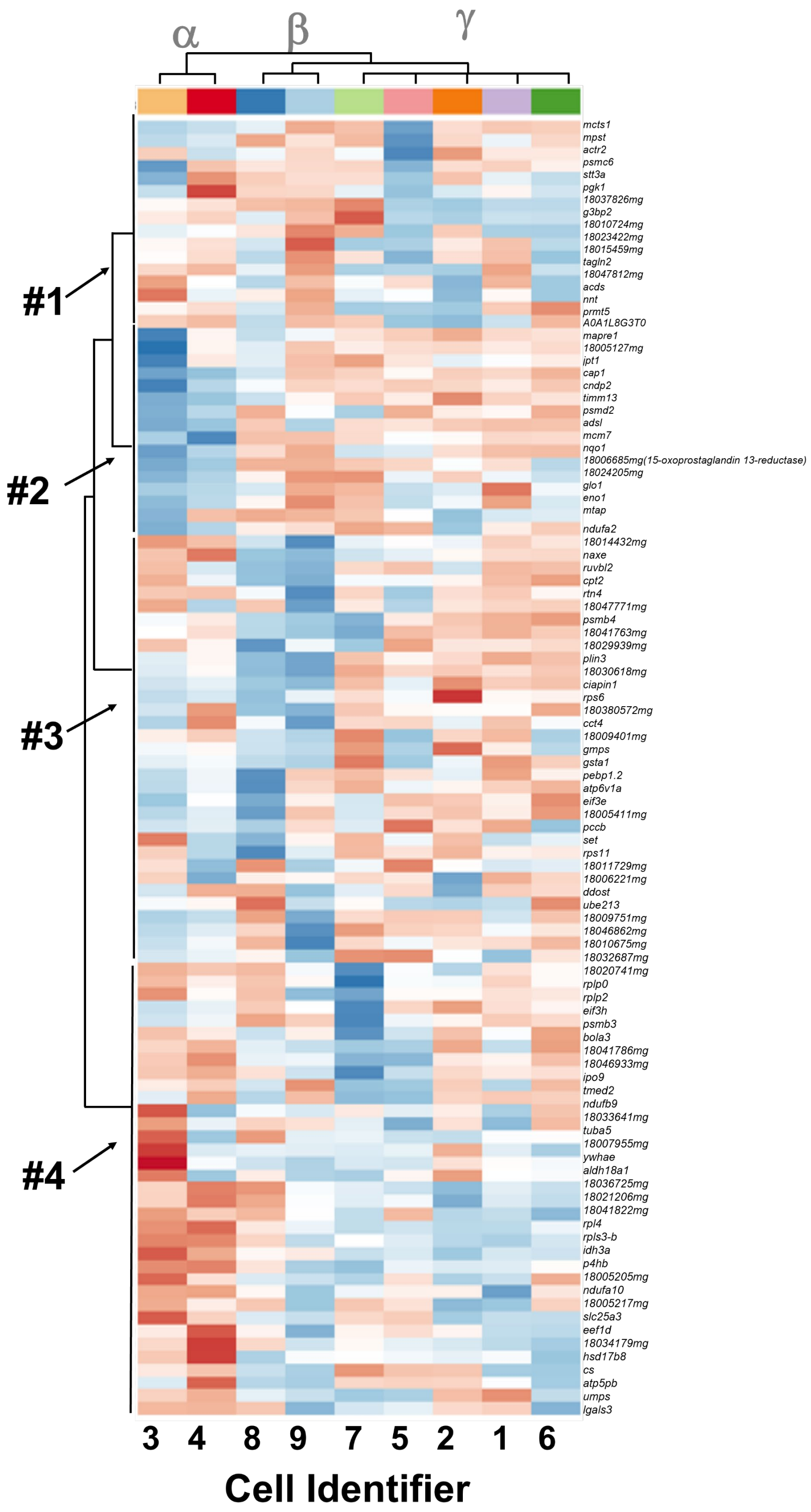

**Figure S3.** Close-up of the hierarchical cluster analysis (HCA)–heatmap (**Fig. 5D**). The 100 most significantly differently abundant proteins are shown. Each protein is labeled by the name of the corresponding gene following the *Xenopus* nomenclature.
